# Genomic basis of multidrug-resistance, mating, and virulence in *Candida auris* and related emerging species

**DOI:** 10.1101/299917

**Authors:** José F. Muñoz, Lalitha Gade, Nancy A. Chow, Vladimir N. Loparev, Phalasy Juieng, Elizabeth L. Berkow, Rhys A. Farrer, Anastasia P. Litvintseva, Christina A. Cuomo

**Affiliations:** Broad Institute of MIT and Harvard, Cambridge, MA USA; Mycotic Diseases Branch, Centers for Disease Control and Prevention, Atlanta, GA USA; Biotechnology Core Facility Branch, Centers for Disease Control and Prevention, Atlanta, GA USA

## Abstract

*Candida auris* is an emergent fungal pathogen of rising public health concern due to increasing reports of outbreaks in healthcare settings and resistance to multiple classes of antifungal drugs. While distantly related to the more common pathogens *C. albicans* and *C. glabrata*, *C. auris* is closely related to three rarely observed and often multidrug-resistant species, *C. haemulonii, C. duobushaemulonii and C. pseudohaemulonii*. Here, we generated and analyzed near complete genome assemblies and RNA-Seq-guided gene predictions for isolates from each of the four major *C. auris* clades and for *C. haemulonii, C. duobushaemulonii and C. pseudohaemulonii*. Our analyses mapped seven chromosomes and revealed chromosomal rearrangements between *C. auris* clades and related species. We found conservation of genes involved in mating and meiosis and identified both *MTL***a** and *MTL*α *C. auris* isolates, suggesting the potential for mating between clades. Gene conservation analysis highlighted that many genes linked to drug resistance and virulence in other pathogenic *Candida* species are conserved in *C. auris* and related species including expanded families of transporters and lipases, as well as mutations and copy number variants in *ERG11* that confer drug resistance. In addition, we found genetic features of the emerging species that likely underlie differences in virulence and drug response between these and other *Candida* species, including genes involved in cell wall structure. To begin to characterize the species-specific genes important for antifungal response, we profiled the gene expression of *C. auris* in response to voriconazole and amphotericin B and found induction of several transporters and metabolic regulators that may play a role in drug resistance. This study provides a comprehensive view of the genomic basis of drug resistance, potential for mating, and virulence in this emerging fungal clade.

## Introduction

*Candida auris* is an emerging fungal pathogen of increasing concern due to high drug resistance and high mortality rates^1,2^. In addition, outbreaks have been reported in hospital settings, suggesting healthcare transmission^2,3^. *C. auris* clinical isolates are typically multidrug-resistant (MDR), with common resistance to fluconazole and variable susceptibility to other azoles, amphotericin B, and echinocandins^2^. *C. auris* causes bloodstream and other invasive and superficial infections, similar to a group of rarely observed, phylogenetically related species including *C. haemulonii, C. duobushaemulonii and C. pseudohaemulonii*^4,5^. These species also display MDR, most commonly to amphotericin B and also reduced susceptibility to azoles and echinocandins^4,5^. Together, *C. auris* and these closely related species represent an emerging clade of invasive fungal pathogens, which are not only difficult to treat, but also difficult to identify using standard laboratory methods^6^, generally requiring molecular methods for proper identification.

Initial whole genome analysis of *C. auris* isolates from Pakistan, India, South Africa, Japan and Venezuela identified four clades that are specific to each geographic region, suggesting that each of these clades emerged nearly simultaneously in different regions of the world^2^. Including data from other recent studies, the current representation of each clade is as follows: clade I comprises isolates from India, Pakistan and the United Kingdom^2,3,7^, clade II from Japan and South Korea^2,8^, clade III from South Africa, and clade IV from Venezuela^2^. Analyses of SNPs identified from whole genome sequence^2,9^ and of multilocus sequence typing (MLST)^10,11^ found low genetic diversity between isolates within each *C. auris* clade. Little is known about whether phenotypic differences exist among isolates from different *C. auris* clades; one noted difference is that isolates from India and South Africa are capable of assimilating N-acetylglucosamine while isolates from Japan and South Korea were not able to process this compound^2^.

Phylogenetic studies revealed that *Candida auris* belongs to the *C. haemulonii* clade and is distantly related to the more common human pathogenic species including *C*. *albicans* and *C*. *glabrata* ^4,5,12,13^. However, these prior studies have not clearly resolved the relationships of species within the *C. haemulonii* clade, including *C. haemulonii, C. duobushaemulonii and C. pseudohaemulonii*^4,5,12,13^. *C. lusitaniae*, a rarely observed cause of infection^14^, is a sister clade to this group of emerging species^13,15^. Despite the fact that *C. auris* is highly divergent from other Saccharomycetales yeasts from the CTG clade, which includes the common human pathogens *C. albicans*, *C. tropicalis* and *C. parapsilosis,* most of our limited current knowledge of *C. auris* resistance and virulence had been inferred based on conservation of genes associated with drug resistance and virulence in *C. albicans* or in *C. glabrata*, which is part of the distantly related *Nakaseomyces* clade. Initial comparative analysis of the gene content of one isolate of *C. auris* with *C. albicans* found that some orthologs associated with antifungal resistance are present in *C. auris*, including drug transporters, secreted proteases and manosyl transferases^13^. In addition, preliminary genomic studies showed that the targets of several classes of antifungal drugs are conserved in *C. auris*, including the azole target lanosterol 14 α-demethylase (*ERG11*), the echinocandin target 1,3-beta-glucan synthase (*FKS1*), and the flucytosine target uracil phosphoribosyl-transferase (*FUR1*)^2,16^. Furthermore, point mutations associated with drug resistance in other species are observed in many clinical isolates and are associated with *C. auris* clades^2,7,16^. Recently, two *ERG11* mutations (Y132F and K143R) in *C. auris* were found to increase resistance to fluconazole^17^, however the role of other genes in drug resistance and virulence has not been reported for this group of emerging MDR species.

While efforts to sequence *C. auris* genome provided an initial view of genome content^9,13^, available assemblies in GenBank are highly fragmented, inconsistently annotated, and do not provide a complete representation of all *C. auris* clades. The related species *C. haemulonii* and *C. duobushaemulonii* were recently sequenced ^18,19^. Here, we generated and annotated highly complete genome assemblies for isolates from each of the clades of *C. auris* as well as for the related species *C. haemulonii*, *C. duobushaemulonii* and *C. pseudohaemulonii*. Comparison of these genomes to other sequenced *Candida* species revealed that *C. auris* has notable expansions of genes linked to drug resistance and virulence in *C. albicans*, including families of oligopeptide transporters, siderophore-based iron transporters, and secreted lipases. We used RNA-Seq to examine the response of two isolates of *C. auris* to antifungal drugs and detected the upregulation of transporters and metabolic regulators that have been previously associated with drug resistance in *C. albicans*. In addition, we also observed several genes that were either unique or expanded in *C. auris* and related species were upregulated in response to azole and amphotericin B exposure. We also found evidence that *C. auris* may be capable of mating and meiosis, based on the identification of both mating types in the populations, conservation of genes involved in mating and meiosis, and detection of chromosomal rearrangements between two clades. These results revealed fundamental insights into the evolution of drug resistance and pathogenesis in *C. auris* and closely related species and the potential for mating and recombination between the *C. auris* clades.

## Results

### Genome characteristics of *Candida auris* and closely related species

We generated highly complete genome assemblies for four *Candida auris* isolates and for three phylogenetically related species, *C. haemulonii*, *C. duobushaemulonii,* and *C. pseudohaemulonii.* Most of these isolates displayed increased resistance to fluconazole and also increased resistance to voriconazole or amphotericin B (**Table S1**). The *Candida auris* assemblies represent each of the four major clades^2^, including an updated assembly for strain B8441 (clade I) and the first representatives of clade II (strain B11220), clade III (strain B11221) and clade IV (strain B11243). As *C. auris* is distantly related to other previously sequenced *Candida* species, we also sequenced genomes of the closely related species *C. haemulonii* (strain B11899), *C. duobushaemulonii* (strain B09383), and *C. pseudohaemulonii* (strain B12108) to enable comparative genomic analysis. The genome assemblies of *C. auris* (B8441 and B11221), *C. haemulonii*, and *C. duobushaemulonii* were sequenced using PacBio and Illumina, whereas the other two *C. auris* strains and *C. pseudohaemulonii* were sequenced only with Illumina (**Table 1**; **Methods**). The genome assemblies of *C. auris* range from 12.1 Mb in B11220 to 12.7 Mb in B11221. The *C. auris* B8441 and B11221 genome assemblies were organized in 15 and 20 scaffolds, respectively, of which 7 scaffolds included most of the sequenced bases in both strains (**Table 1**). This represents an improvement upon the previously generated genome assembly of *C. auris* strain 6684 (clade I)^13^, which consists of 99 scaffolds (759 contigs) that include contig gaps and has an inflated number of predicted genes, based on our analysis (see below; **Methods**). The genome assemblies of Illumina data for *C. auris* B11220 (clade II) and B11243 (clade IV) are less contiguous than the assemblies that included long reads, consisting of 324 or 285 contigs, respectively (**Table 1**). The genome assemblies of *C. haemulonii*, *C. duobushaemulonii,* and *C. pseudohaemulonii* were also highly contiguous with 11, 7 and 36 scaffolds, respectively (**Table 1**). Comparison of these four *C. auris* genomes and those of the three related species revealed that the genome sizes are very similar, ranging in size from 12.1 Mb in *C. auris* B11220 to 13.3 Mb in *C. haemulonii*, similar to that of *C. lusitaniae* (12.1 Mb) and other *Candida* species (**Table S2**).

**Table 1.** Genome assembly statistics of *Candida auris* and closely related species^*^

| Species | *Candida auris* |  |  |  | *C. hae* | *C. duo* | *C. pseu* |
| --- | --- | --- | --- | --- | --- | --- | --- |
| Strain | B8441 | B11221 | B11220 | B11243 | B11899 | B09383 | B12108 |
| Clade | I | III | II | IV | . | . | . |
| Country of origin | Pakistan | South Africa | Japan | Venezuela | Israel | USA | Venezuela |
| Total assembly size (Mb) | 12.4 | 12.7 | 12.1 | 12.3 | 13.3 | 12.3 | 12.6 |
| Chromosomes | 7 | 7 | . | . | . | . | . |
| Assembly anchored (%) | 98.8 | 97.2 | . | . | . | . | . |
| Scaffolds | 15 | 20 | 324 | 240 | 11 | 7 | 36 |
| Contigs | 18 | 23 | 324 | 265 | 11 | 7 | 41 |
| Scaffold N50 (Mb) | 1.1 | 2.4 | 0.06 | 0.09 | 1.7 | 3.3 | 0.64 |
| Scaffold N90 (kb) | 777 | 949 | 19.5 | 27.1 | 952 | 788 | 227 |
| GC content (%) | 45.2 | 45.3 | 45.0 | 45.0 | 45.3 | 46.9 | 47.2 |
| Protein coding genes | 5,421 | 5,527 | 5,546 | 5,601 | 5,410 | 5,331 | 5,288 |

\**C. hae* = *C. haemulonii*; *C. duo* = *C. duobushaemulonii*; *C. pseu* = *C. pseudohaemulonii*

The assemblies of the *C. auris* clades are highly identical, sharing an average pairwise nucleotide identity of 98.7% across all four clades (I, II, III and IV); clades II and III appear the most similar and share 99.3% identity. By contrast, comparison of nucleotide genome identity between species highlights greater interspecies genetic divergence: 88% between *C. auris* and each of the three related species (*C. haemulonii*, *C. duobushaemulonii,* and *C. pseudohaemulonii*), 89% between *C. haemulonii* and *C. duobushaemulonii*, and 92% between *C. duobushaemulonii* and *C. pseudohaemulonii*. To provide a quantitative measure of the degree of genetic variation within the *C. auris* population, we calculated the genome-wide nucleotide diversity (π) using SNPs identified from the isolate sequences of Lockhart *et al*^2^. The estimated π is lowest for *C. auris* clade I 0.00050, and slightly higher for clades III and IV (0.00079 and 0.00075, respectively); π for all *C. auris* isolates is 0.0039, ~17-fold higher than the intra-clade levels. Comparing to other human fungal pathogens, this level of diversity is lower than that reported in *C. neoformans* var. *grubii*, (π = 0.0074, ^20^), but higher than the extreme clonality reported for *Trichophyton rubrum* (π = 0.00054, ^21^). These comparisons support that the different clades of *C. auris* comprise a single species well separated from the related MDR species.

As an independent assessment of the genome assembly size and structure, we generated optical maps of the *C. auris* B8441 and B11221 isolates (**Figure S1**). Consistent with the assemblies of these isolates, the maps had seven linkage groups; nearly all of the genome assemblies were anchored to the optical maps (98.8 % of B8441 and 97.2 % of B11221; **Figure S1; Table 1**). This supports the presence of seven chromosomes in *C. auris*, consistent with the chromosome number found in previous studies using electrophoretic karyotyping by pulsed-field gel electrophoresis (PFGE)^22^. While the genomes of *C. auris* are highly syntenic, we found evidence of a few large chromosomal rearrangements between *C. auris* B8441 and B11221 based on comparison of the assemblies and the optical maps (**Figures 1**, **S1**). We confirmed that the junctions of these rearrangements are well supported in each assembly; these regions show no variation in the depth of PacBio and Illumina aligned reads and there is no evidence of assembly errors across these rearrangement breakpoints. We additionally independently identified structural variants based on the read alignments to the B8441 and B11221 assemblies and recovered each of the rearrangements present in these assemblies. These large chromosomal rearrangements included one inversion of 136 kb between B8441 sc01 and B11221 sc01, a 274 kb translocation between B8441 sc08 and B11221 sc03, and a 300 kb translocation between B8441 sc10 and B11221 sc01 (**Figure 1**). The genomes of *C. auris*, *C. haemulonii*, *C. duobushaemulonii,* and *C. pseudohaemulonii* showed limited chromosomal rearrangements between each other, mostly intra-chromosomal inversions between *C. auris* and *C. haemulonii, C. duobushaemulonii,* and *C. pseudohaemulonii,* and large chromosomal translocations between *C. haemulonii*, *C. duobushaemulonii,* and *C. pseudohaemulonii* (**Figure 1a**). These rearrangements between *C. auris* clades, and between species, could potentially prevent genetic exchange between these groups, since some crossover events will generate missing chromosomal regions or other aneuploidies and may result in nonviable progeny.

**Figure 1.**
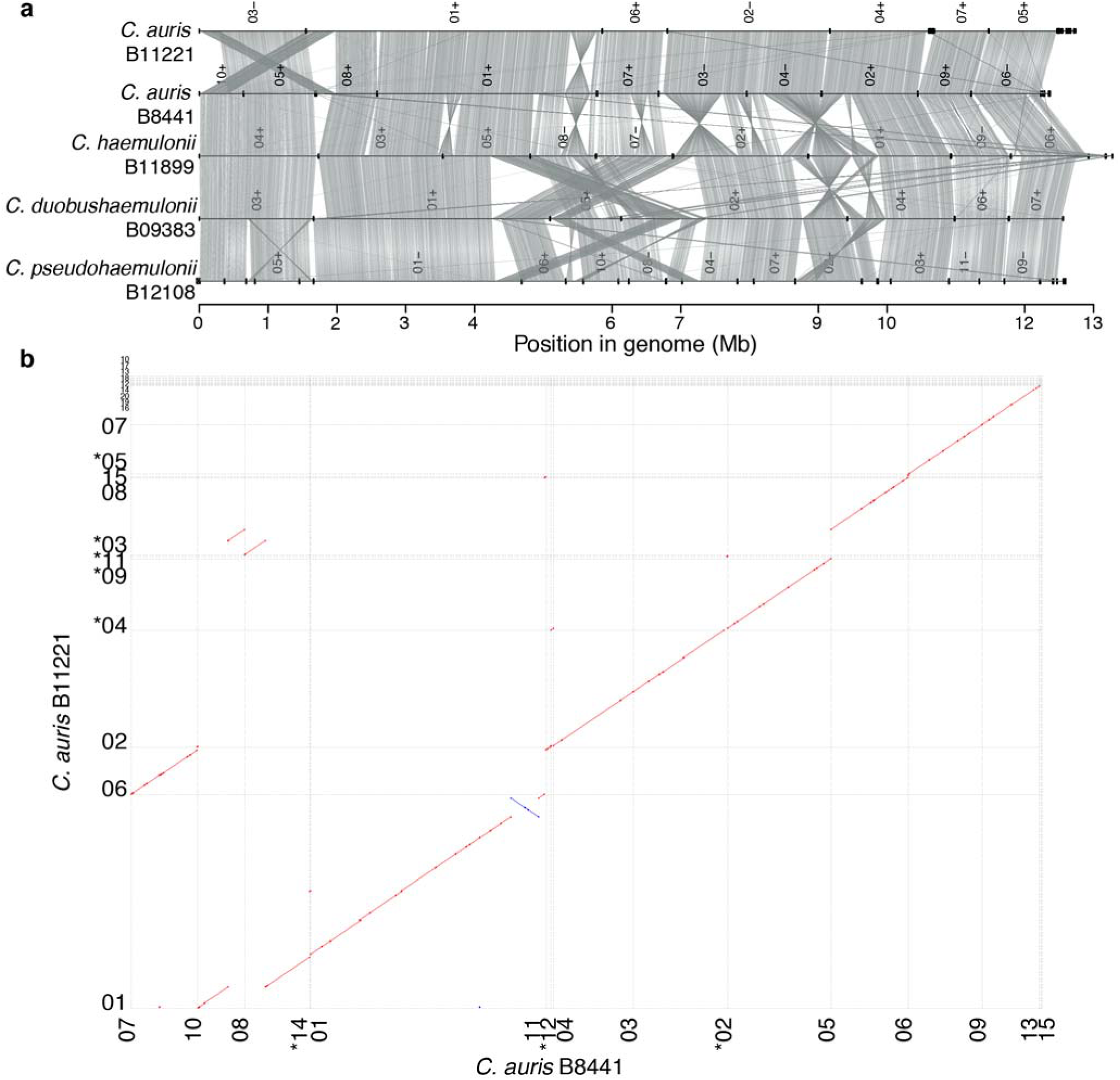
Whole genome conservation, structure and synteny. **(a)** Genome wide gene synteny among *Candida auris*, *C. haemulonii*, *C. duobushaemulonii*, and *C. pseudohaemulonii*. To determine synteny regions of conserved gene order we identified orthologs between these isolates and then plotted chains of orthologs (grey lines). Isolate names are shown to the left of their genomes, which are represented by horizontal black lines, with vertical lines indicating scaffold borders, and their identifiers listed above (+/− = orientation). **(b)** Shared synteny regions based on whole genome alignments between *C. auris* B8441 and B11221.

### Evolution of mating-type locus in the emerging multidrug-resistant species including *C. auris*

We characterized the mating type locus in *C. auris* and identified representatives of both mating types. The mating locus structure is highly conserved compared to closely related species including *C. lusitaniae* and other Saccharomycetales yeasts from the CTG clade *Candida* (**Figure 2**). Many *Candida* species, including diploid asexual species, have heterothallic *MTL* idiomorphs, and mating occurs between cells of opposite mating type, *MTL***a** and *MTL*α^23^. We found that the genes flanking the *MTL* locus in some species from the CTG clade *Candida*, namely the phosphatidylinositol kinase gene (*PIK1*), the oxysterol binding protein gene (*OBP1*) and the poly(A) polymerase gene (*PAP1*) were adjacent in all sequenced genomes of the emerging MDR clade (**Figure 2a**). Previous work reported that the genome of *C. auris* 6684 included the *MTL* flanking genes but did not identify either *MTL***a** or *MTL*α genes at this locus^13^. In the chromosome level genome assemblies of *C. auris* (B8441 and B11221), the *PIK1/OBP1/PAP1* genes are present at a single locus in chromosome 3 in B8441 (sc05) and B11221 (sc03). Notably, we found all *C. auris* isolates contained either the *MTL***a** and *MTL*α idiomorphs at this locus and that gene order was conserved compared to the *C. lusitaniae MTL*^24^ (**Figure 2a**; **Table S3**). We found that the *MTL***a** is present in *C. auris* B8441 and 6684 (clade I), B11243 (clade IV), and *C. pseudohaemulonii*, spanning 14.9 kb (**Figure 2a**). By contrast, *C. auris* B11220 (clade II), B11221 (clade III), and *C. haemulonii* and *C. duobushaemulonii* contain *MTL*α, spanning 14.3 kb (**Figure 2a**). Phylogenetic analysis of the non-mating flanking genes (*PIK1/OBP1/PAP1*) supports the inheritance of idiomorphs of these genes with the *MTL**a*** and *MTL*α genes (**Figure 2b**). Upon further examination and manual annotation, we determined that **a**1*/***a**2 are present in *MTL***a** isolates, and α1 is present in *MTL*α isolates; as in *C. lusitaniae*, the MTL α locus is missing the α2 gene^24^ (**Figure 2a**). RNA-Seq data was used to guide gene prediction of **a**1 and **a**2, and further established that both genes are expressed in B8441 (**Figure S2**), supporting the hypothesis that these genes could be functional, and that the MTL locus could be used to classify *C. auris* isolates. To further characterize the evolution of mating type in the population, we examined the *MTL* locus using 50 isolates from Lockhart *et al*.^2^ and found that all isolates from clade I and IV have *MTL***a**, whereas all isolates from clade II and III had *MTL*α (**Figures 2c**, **S3**). The fact both **a** and α *MTL* alleles are present and expressed in *C. auris* suggests this species may be capable of mating, either in the ancestral population of the outbreak clades or if the expanding outbreak results in the presence of isolates from both mating types in a geographic area.

**Figure 2.**
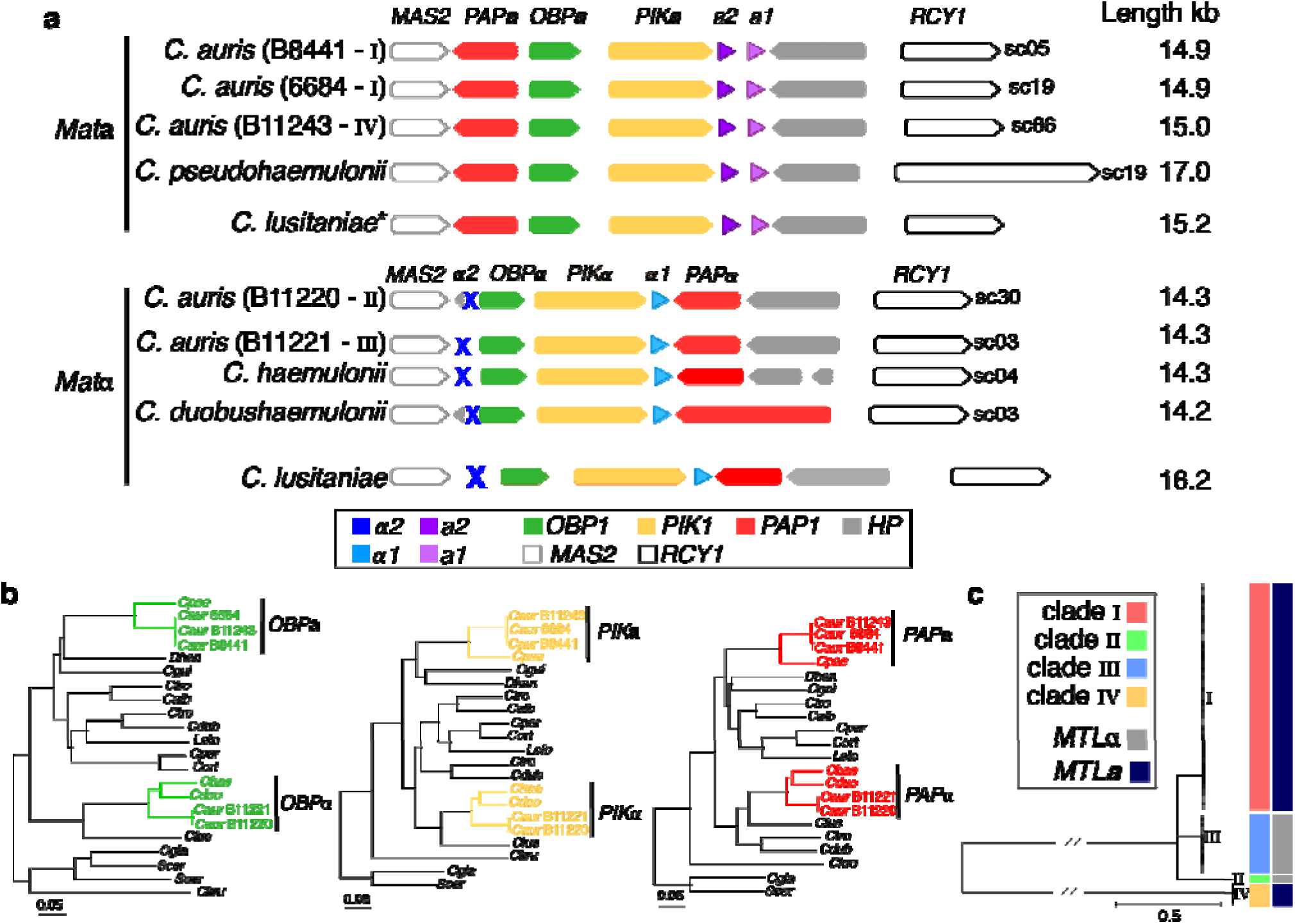
Identification of Mating-type loci (*MTL*) in *Candida auris* and closely related species. **(a)** Synteny schema depicting orientation and conservation of the color-coded *MTL* idiomorphs and genes adjacent to the *MTL*. The putative MTL in *C. auris*, *C. haemulonii*, *C. duobushaemulonii*, and *C. pseudohaemulonii* are shown in comparison with the *MTL**a*** and *MTL*α idiomorphs from *C. lusitaniae*. **(b)** Phylogenetic analysis of the non-mating flanking genes (*PIK1/OBP1/PAP1*) showing the inheritance of idiomorphs of these genes within the *MTL***a** and *MTL*α loci. Branch lengths indicate the mean number of changes per site. **(c)** Phylogenetic tree of *C. auris* isolates from Lockhart et al. 2016. Isolates are color-coded according the clades (I, II, III and IV) and mating type (*MTL***a** and *MTL*α). Figure S3 shows isolates, origin and the normalized depth read coverage of mapped positions for all isolates aligned to B8441 (*MTL***a**) and B11221 (*MTL*α), supporting the classification into *MTL***a** or *MTL*α.

We examined the conservation of genes involved in meiosis to provide additional support for potential mating in *C. auris*. We found that many of the key meiotic genes are similarly conserved between *C. lusitaniae*, *C. guilliermondii*, *C. auris*, *C. haemulonii*, *C. duobushaemulonii* and *C. pseudohaemulonii* (**Table S3**). Some genes involved in meiosis in *S. cerevisiae* appeared absent in these species including the recombinase *DMC1* and cofactors (*MEI5* and *SAE3*), synaptonemal-complex proteins (*ZIP1* and *HOP1*), and genes involved in crossover interference (*MSH4* and *MSH5*; **Table S3**). A small number of genes involved in meiosis were present in *C. lusitaniae* but absent in *C. auris*, *C. haemulonii*, *C. duobushaemulonii,* and *C. pseudohaemulonii*. In *C. auris*, the DNA recombination and repair genes *RAD55* and *RAD57* appear to be absent, however *RAD55* is widely absent in the CTG *Candida* clade while the *RAD51* paralog of the *RAD55*/*RAD57* complex is present. *C. lusitaniae*, despite the loss of many of the same meiotic genes, undergoes meiosis during sexual reproduction involving diploid intermediates^24^. The fact that most components of the mating and meiosis pathways are similarly conserved in *C. auris* and closely related species including *C. lusitaniae* suggests these species may have the ability to mate and undergo meiosis as observed for *C. lusitaniae*.

### Phylogenetic position of *C. auris*, *C. haemulonii*, *C. duobushaemulonii,* and *C. pseudohaemulonii*

Using the complete genomes, we estimated a strongly supported phylogeny of *C. auris*, *C. haemulonii*, *C. duobushaemulonii,* and *C. pseudohaemulonii*, relative to other species from the order Saccharomycetales including *C. lusitaniae*, *C. tropicalis*, *C. albicans* and *C. glabrata* (**Table S2**). Based on a concatenated alignment of 1,570 single copy core genes, a well supported maximum likelihood tree placed *C. auris*, *C. haemulonii*, *C. duobushaemulonii,* and *C. pseudohaemulonii* as a single clade, confirming the close relationship of these species (100% of bootstrap replicates; **Figure 3a**). The *C. auris* clades appear more recently diverged based on short branch lengths within this species. Previous phylogenetic analyses had shown conflicting relationships between *C. auris*, *C. haemulonii*, *C. duobushaemulonii,* and *C. pseudohaemulonii*^4,5,12,13^. Our phylogenetic analysis strongly supports that *C. duobushaemulonii* and *C. pseudohaemulonii* are most closely related to each other, and form a sister group to *C. haemulonii*, which appeared as the more basally branching species (**Figure 3a**). The most closely related species to this MDR clade is *C. lusitaniae*, which is the more basally branching member of this group (**Figure 3a**).

**Figure 3.**
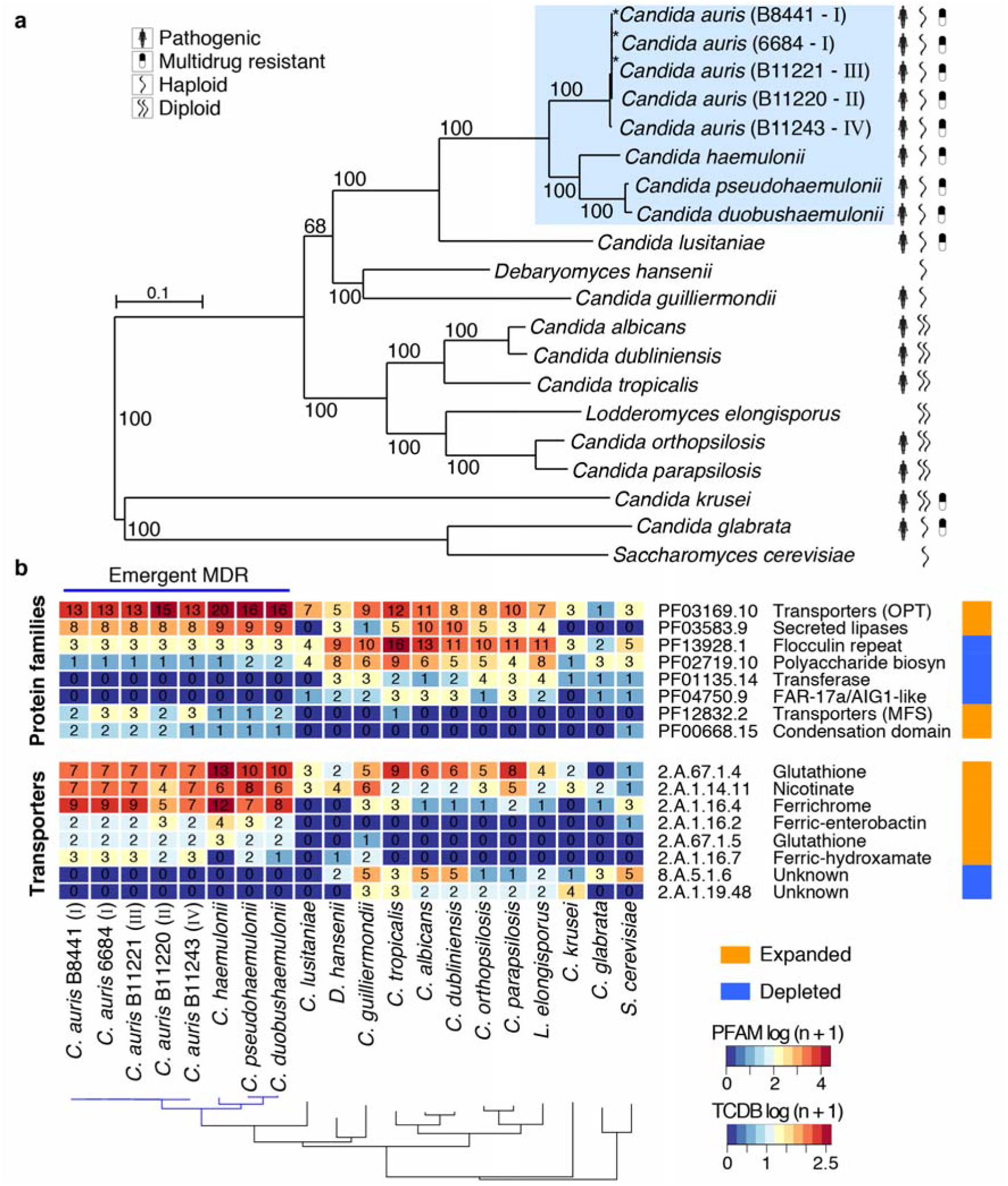
Phylogenomic and gene family changes across *Candida auris* and closely related species. **(a)** Maximum likelihood phylogeny using 1,570 core genes based on 1,000 replicates, among 20 annotated genome assemblies, including *Candida auris, C. haemulonii* (B11899), *C. duobushaemulonii* (B09383), and *C. pseudohaemulonii* (B12108), and closely related species. Branch lengths indicate the mean number of changes per site. **(b)** Heatmap depicting results of protein family enrichment analysis (PFAM domains; corrected *p-value* < 0.05) comparing the gene content of *C. auris* strains representing each clade, *C. haemulonii*, *C. duobushaemulonii*, and *C. pseudohaemulonii*, and other closely related species, including *C. lusitaniae*, *C. albicans*, *C. krusei* and *C. glabrata*. Values are colored along a blue (low counts) to red (high counts) color scale, with color scaling relative to the low and high values of each row. Each protein family domain has a color code (right) indicating whether expanded or depleted.

### Gene family expansions supported mechanisms of drug resistance and virulence

Gene annotation of the *C. auris* genomes was performed using RNA-Seq paired-end reads to improve gene structure predictions (**Methods**). The predicted gene number was highly similar across all *C. auris* genomes as well as in *C. haemulonii*, *C. duobushaemulonii,* and *C. pseudohaemulonii*. In *C. auris*, the number of protein-coding genes varied between 5,421 in B8441 and 5,601 in B11243. For *C. haemulonii*, *C. duobushaemulonii,* and *C. pseudohaemulonii* the numbers were very similar, ranging from 5,288 to 5,410 predicted genes (**Table 1**; **Figure S4a**). High representation of core eukaryotic genes provides evidence that these genomes are nearly complete; 96-98% of these conserved genes are found in all annotated genome assemblies (**Figure S4b**). By examining orthologous genes in *C. auris*, *C. haemulonii*, *C. duobushaemulonii, C. pseudohaemulonii*, and twelve additional Saccharomycetales genomes, including *C. lusitaniae*, *C. tropicalis*, *C. albicans* and *C. glabrata*, we found a total of 2,379 core ortholog clusters had representative genes from all twenty analyzed genomes (**Figure S4**). We found a small number of unique genes in the *C. auris* clades ranging from 15 (B8441; clade I) to 54 (B11221; clade III) genes, and higher numbers in the three related species (203 in *C. haemulonii*, 83 in *C. duobushaemulonii,* and 88 *C. pseudohaemulonii*) (**Figure S5**). The unique genes in *C. auris* clades include oligopeptide and ABC transporters (**Table S4**). While unique glycophosphatidylinositol (GPI)-anchored proteins were identified in *C. auris*, we found high conservation across *C. auris*, *C. haemulonii*, *C. duobushaemulonii, C. pseudohaemulonii* of seven (GPI)-anchored proteins (*PLB3*, *IFF4*, *PGA52*, *PGA26*, *CSA1*, *HYR3* and *PGA7*) that were upregulated during *C. auris* biofilm formation and are associated with antifungal resistance and biofilm mechanisms in *C. albicans*^25^(**Table S4**).

To characterize changes in gene content that may play a role in the evolution of multidrug-resistance and virulence in *C. auris*, *C. haemulonii*, *C. duobushaemulonii,* and *C. pseudohaemulonii*, we searched for expansions or contractions in functionally classified genes compared to other related species (**Table S2**). We identified PFAM domains that were significantly enriched or depleted (**Methods**; **Figures 3b, S4**). Domains associated with transmembrane transporters (OPT, MFS) and secreted lipases (LIP) were enriched in *C. auris*, *C. haemulonii*, *C. duobushaemulonii,* and *C. pseudohaemulonii* compared to other genomes (*q-* value < 0.05, Fisher’s exact test; **Figure 3b**). We therefore further classified transmembrane transporters using the Transporter Classification Database (TCDB) and found that the higher copy number of transporters in the emergent MDR clade could be attributed to oligopeptide transporters (*OPT*) and siderophore iron transporters (*SIT*). In *C. albicans*, *OPT* transporters enable uptake of small peptides; the expression of some *OPT* transporters are up-regulated by azole drugs^26,27^. While most of the 14 *OPT* genes found in *C. auris* had orthologs in *C. albicans* (*OPT1-8*), a subset of these transporters appear recently duplicated in the MDR emergent clade, including an expansion of three *OPT1*-like transporters, and five transporters similar to *OPT2*, *OPT3*, and *OPT4* (**Figures 4a**, **S6**). Most of the *OPT* genes (up to 8) are located in a conserved locus among emerging MDR species encompassing 296 kb of chromosome 6; this gene family appears to have expanded by tandem duplication (**Figures 4c**). In *C. albicans*, iron transporters include the siderophore transporter *SIT1* and the iron permeases *FTR1* and *FTR2*; a subset of these transporters is uniquely expanded in *C. auris* and closely related species, including the expansion of 14 ortholog groups in *C. auris* related to *C. albicans SIT1* (**Figures 4b**, **S6**). Secreted lipases are also expanded in the genomes of *C. auris* and closely related species (*q-*value < 0.05, Fisher’s exact test; **Figure 3b**). The *C. auris* clade has similar counts of lipases relative to *C. albicans* and *C. dubliniensis*, however, these proteins are expanded relative to more closely related human pathogenic species, including *C. lusitaniae*, *C. guilliermondii*, *C. krusei*, and *C. glabrata* (**Figure 3b**). Phylogenetic analysis suggested independent evolutionary trajectories of secreted lipases in emerging *C. auris* and related species, where the most recent ortholog family of lipases includes *C. albicans LIP4*, *5*, *8* and *9* (**Figure S7**). The secretion of lipases may be important during infection for nutrient acquisition, adaptation, virulence and immune evasion^28^; we identified a predicted secretion signal in all lipases encoded by *C. auris* and related species, supporting an extracellular role in these emergent MDR species.

**Figure 4.**
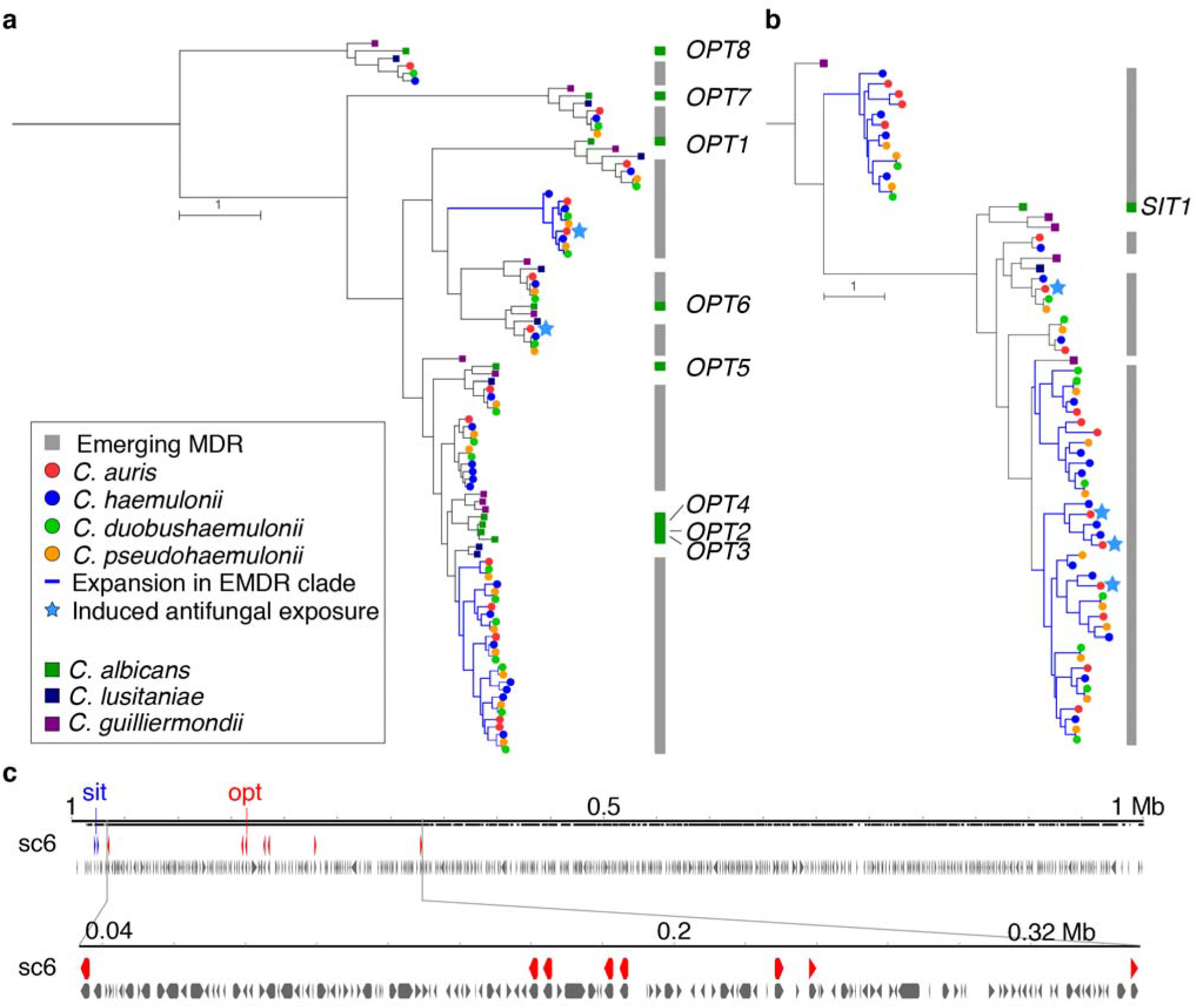
Phylogenetic relationships of oligopeptide transporters (*OPT*) and siderophore iron transporters (*SIT*) families. Phylogenetic trees estimated by maximum likelihood with RAxML showing expansion of *OPT* **(a)** and *SIT* **(b)** transporter families in *C. auris* and *C. haemulonii*, *C. duobushaemulonii* and *C. pseudohaemulonii*. Each species has a color code and lineage-specific expansions (blue branches) can be seen in *C. auris* and closely related species relative to the close ancestor *C. lusitaniae* and *C. albicans*. Orthologs of *OPT* and *SIT* transporters in *C. albicans* are depicted alongside each tree. **(c)** Chromosome view depicting genes and orientation located in chromosome 6 (B8441 scaffold05). This region highlights expansion and tandem duplication of eight *OPT* class transporters (in red).

### Conservation of known drug resistance and pathogenesis-associated genes

The isolates selected for genome sequencing display increased resistance to antifungal drugs. Increased resistance to fluconazole is most commonly observed in *C. auris* with some isolates also displaying increased resistance to voriconazole or amphotericin B (**Table S1;**^2^). The three related species all displayed increased amphotericin B resistance, and two were also resistant to fluconazole. All isolates appeared sensitive to the echinocandins tested, anidulafungin and caspofungin (**Table S1**). Most of the genes associated with drug resistance and pathogenesis in *C. albicans* are conserved in *C. auris*, *C. haemulonii*, *C. duobushaemulonii,* and *C. pseudohaemulonii*. We identified orthologs of genes noted to confer drug resistance in *C. albicans*, either by acquiring point mutations, increasing transcription, or copy number variation. The annotated genome assemblies of *C. auris*, *C. haemulonii*, *C. duobushaemulonii,* and *C. pseudohaemulonii* contain a single copy of the *ERG11* azole target and the *UPC2* transcription factor that regulates expression of genes in the ergosterol pathway, as well as all the gene components of the ergosterol biosynthesis pathway (**Table S5**). Several of the sites in *ERG11* subject to drug resistant mutations in *C. albicans* are similarly mutated in drug resistant *C. auris* isolates (at positions Y132, K143, and F126, as reported previously^2^). Analysis of the annotated genome assemblies of *C. auris* isolates agrees with prior SNP analysis^2^, with the exception of F126L mutation in B11221 (clade III), initially reported as F126T for this isolate though this was recently corrected to F126L^29^. The F126L mutation is also observed in a separate genomic study of *C. auris*^7^ and in drug resistant *C. albicans*^30^ (**Figure S8**). The Y132F mutation observed in two *C. auris* clades is also found in *C. pseudohaemulonii*, suggesting this site may contribute to the increased azole resistance of this *C. pseudohaemulonii* isolate (**Table S1**). Other sites in *ERG11* that confer drug resistance in *C. albicans* do not display drug resistant mutations in *C. auris*, *C. haemulonii*, *C. duobushaemulonii,* and *C. pseudohaemulonii* (**Figure S8**). In addition, we found that not all *C. auris* isolates have a single copy of *ERG11*. Using copy number variation (CNV) analysis of the Illumina read depth of 47 previously sequenced isolates, we found a total of 6 large duplicated regions (CNVnator *p*-value < 0.01) ranging in size from 12 to 153 kb (**Table S6, Methods**). The largest duplicated region of 153 kb is present in two isolates (B11227 and B11229) from clade III and encompassed 62 genes including *ERG11* (**Figure S9**; **Table S6**). These two isolates also display increased resistance to fluconazole (**Table S1**). Long read assemblies of these isolates could be used to examine the chromosomal context of this duplication.

Additionally, we identified orthologs of transporters from the ATP binding cassette (*ABC*) and major facilitator superfamily (*MFS*) classes of efflux proteins that are involved in clinical antifungal resistance in *C. albicans* by the overexpression of ABC transporter family *CDRs* (*CDR1* and *CDR2*) and the MFS transporter *MDR1* ^31,32^. We identified a single copy of the multidrug efflux pump *MDR1* in all sequenced isolates (**Table 2**). We further characterized presence of *CDR* genes and *MDR1* in the *C. auris* population by examining CNV and gene conservation across the 47 isolates from Lockhart *et al*.^2^, and did not detect any copy number variation of these transporters across this set (**Table S7**). Candidate multidrug transporters similar to *CDR1*, *CDR2*, and related genes include 5 genes in most genomes (*C. auris* B8441, B11220, and B11243, *C. haemulonii*, and *C. pseudohaemulonii*), 6 genes in *C. auris* B11221, and 4 genes in *C. duobushaemulonii* (**Table 2**; **Figure S10**). Phylogenetic analysis of these ABC transporters showed that one of the genes in *C. auris* is related to *CDR1*/*CDR2*/*CDR11*, two genes are related to *CDR4*, and two genes are related to *SNQ2*; *C. auris* B11221 has an additional copy of *SNQ2* (**Figure S10**). The *TAC1* transcription factor that regulates expression of *CDR1* and *CDR2* in *C. albicans* is present in two tandem copies in *C. auris*, *C. haemulonii*, *C. duobushaemulonii,* and *C. pseudohaemulonii* (**Table 2**).

**Table 2.** Conservation of genes involved in pathogenesis and drug resistance^*^

| Category | *C. albicans* | *Candida auris* (B8441 - I) | *Candida auris* (B11221 - III) | *Candida auris* (B11220 - II) | *Candida auris* (B11243 - IV) | *Candida auris* (6684 - I) | *Candida haemulonii* | *Candida pseudohaemulonii* | *Candida duobushaemulonii* | *Candida lusitanae* | *Debaryomyces hansenii* | *Candida guilliermondii* | *Candida tropicalis* | *Candida albicans* | *Candida dubliniensis* | *Candida orthopsilosis* | *Candida parapsilosis* | *Lodderomyces elongisporus* | *Candida krusei* | *Candida glabrata* | *Saccharomyces cerevisiae* |
| --- | --- | --- | --- | --- | --- | --- | --- | --- | --- | --- | --- | --- | --- | --- | --- | --- | --- | --- | --- | --- | --- |
| Drug resistance | *ERG11* | 1 | 1 | 1 | 1 | 1 | 1 | 1 | 1 | 1 | 1 | 1 | 1 | 1 | 1 | 1 | 1 | 1 | 1 | 1 | 1 |
|  | *TAC1* | 2 | 2 | 2 | 2 | 2 | 2 | 2 | 2 | 2 | 2 | 1 | 1 | 1 | 1 | 1 | 1 | 1 | 1 | 0 | 0 |
|  | *UPC2* | 1 | 1 | 1 | 1 | 1 | 1 | 1 | 1 | 1 | 1 | 1 | 1 | 1 | 1 | 1 | 1 | 1 | 1 | 2 | 2 |
|  | *MDR1* | 1 | 1 | 1 | 1 | 1 | 1 | 1 | 1 | 1 | 3 | 6 | 2 | 1 | 1 | 3 | 2 | 4 | 0 | 1 | 1 |
|  | *SNQ2*<br>*CDR4*<br>*CDR2*<br>*CDR11*<br>*CDR1* | 5 | 6 | 5 | 5 | 5 | 5 | 5 | 4 | 3 | 6 | 8 | 2 | 5 | 5 | 3 | 6 | 5 | 9 | 5 | 8 |
| Secreted aspartyl proteinases | *SAP1*<br>*SAP2*<br>*SAP8*<br>*SAP3* | 3 | 3 | 2 | 3 | 3 | 2 | 2 | 2 | 2 | 0 | 4 | 3 | 4 | 4 | 3 | 2 | 1 | 0 | 0 | 0 |
|  | *SAP9* | 1 | 1 | 1 | 1 | 1 | 1 | 1 | 1 | 1 | 2 | 1 | 1 | 1 | 1 | 2 | 5 | 1 | 2 | 1 | 3 |
| Secreted lipases | *LIP4, LIP9*<br>*LIP5, LIP8*<br>*LIP2, LIP1*<br>*LIP10, LIP6*<br>*LIP3* | 8 | 8 | 8 | 8 | 8 | 9 | 9 | 9 | 0 | 1 | 1 | 2 | 9 | 9 | 2 | 2 | 1 | 0 | 0 | 0 |
| Cell wall adhesins | *ALS2, ALS5*<br>*ALS1, ALS9*<br>*ALS4, ALS3*<br>*ALS7, ALS6* | 1 | 1 | 1 | 1 | 1 | 2 | 3 | 3 | 1 | 0 | 2 | 4 | 8 | 6 | 1 | 2 | 0 | 0 | 0 | 0 |
|  | *ALS-like* | 2 | 2 | 1 | 2 | 2 | 0 | 1 | 1 | 0 | 0 | 0 | 0 | 0 | 0 | 0 | 0 | 0 | 0 | 0 | 0 |

\* Roman numerals represent *C. auris* clades.

While many gene families involved in pathogenesis in *C. albicans* are present in similar numbers in *C. auris*, *C. haemulonii*, *C. duobushaemulonii,* and *C. pseudohaemulonii*, there are some notable differences such as of cell wall and transmembrane proteins. We identified similar numbers of the secreted aspartyl proteases, lipases and oligopeptide transporters (*OPT*), and only one copy of the *ALS* cell surface family of *C. albicans* (**Table 2**). We also examined whether other genes involved in the *C. albicans* core filamentation response were conserved in the emerging multidrug-resistant species (**Table S4**). While most of these genes are conserved in *C. auris* and closely related species, two genes are absent, candidalysin (*ECE1*) and the hyphal cell wall protein (*HWP1*), both of which are highly expressed in *C. albicans* hyphae^33,34^. Thus, we additionally assessed if any other cell surface families of proteins are enriched in *C. auris* and closely related species. We found a total of 75 genes with a predicted GPI anchor, including genes that were found only in the emerging multidrug resistant clade, including one unique family expanded in *C. auris* (**Table S4**). The most represented protein family domains in these genes included the N-terminal cell wall domain, the aspartyl protease domain, and the fungal specific cysteine rich (CFEM) domain (**Table S4**). The shared profile of these genes across *C. auris* and other MDR species suggests that the more rarely observed species may be similarly primed to become more common human pathogens.

### Transcriptional analysis to voriconazole and amphotericin B in *C. auris* susceptible and resistant strains

To investigate which *C. auris* genes are involved in antifungal resistance, we carried out RNA-Seq of *C. auris* strains B8441 and B11210 (clade I) to profile gene expression changes after exposure to two antifungal drugs, voriconazole (VCZ) and amphotericin B (AMB). B11210 is resistant to AMB and exhibits an elevated MIC to VCZ, while B8441 is susceptible to AMB and displays a low MIC to VCZ (**Table S1**; ^2^). We identified differentially expressed genes (DEGs) in B8441 and B11210 after 2 and 4 hours of drug exposure (**Methods; Table S8; Figure S11a**). The response of the sensitive isolate B8441 to AMB or VCZ involved increased expression of small sets of genes: 39 genes were induced in response to AMB, 21 in response to VCZ, of which 14 were induced by both drugs (fold change (FC) > 2; false discovery rate (FDR) < 0.05; **Figure S11b**). Genes induced upon AMB exposure were enriched in small molecule biosynthetic process and iron homeostasis (enriched GO terms corrected-*P <* 0.05, hypergeometric distribution with Bonferroni correction; **Table S9**). Notably, this set includes genes involved in the *C. albicans* transcriptional response to AMB associated with transport and with lipid, fatty acid, and sterol metabolism^35^ including genes involved in arginine synthesis (*ARG1/ARG3*), ergosterol biosynthesis (*ERG24*), fatty-acid metabolism (*FAS1*/*FAS2*), GPI-linked surface proteins (*PGA7* and *RBT5*), and several iron transporters (class *FTH1* and *SIT1*) (**Table S9; Figure S11c**). Three of these genes (*SIT1*, *PGA7* and *RBT5*) have also been found to be upregulated during *C. auris* biofilm formation^25^, suggesting that cell wall reorganization may be part of the drug response. Genes induced in B8441 in response to both AMB and VCZ were enriched in transmembrane transport and iron transport categories (enriched GO terms corrected-*P <* 0.05, hypergeometric distribution with Bonferroni correction; **Table S8**), including a ferric reductase (*FRP1*), a high affinity iron transporter (*FTH1*), a glucose transporter (*HGT7*), a N-acetylglucosamine transporter (*NGT1*) and an oligopeptide transporter (*OPT1-like*; **Table S9**). Genes only induced with VCZ included the oligopeptide transporter *PTR22.* We also examined expression differences in the resistant isolate B11210. As a large set of genes was identified as differentially expressed compared to the control sample, we focused on the most highly induced or repressed genes (FC > 4; FDR < 0.001; **Figure S11a**). A total of 106 genes were induced in response to AMB and 41 genes in response to VCZ, of which 40 were commonly induced (**Figure S11b**). Genes induced in response to AMB were enriched in transcription and translation processes and in sterol biosynthetic process (enriched GO terms corrected-*P <* 0.05, hypergeometric distribution with Bonferroni correction). Notably, 5 of the 21 genes involved in the ergosterol biosynthesis pathway were highly induced in B11210, including *MVD*, *ERG2*, *ERG1*, *ERG6*, and *ERG13* (**Table S9**; **Figure S11b**; **Figure S12**). This correlates with the maintenance of cell membrane stability, as previously noted for the transcriptional response of *C. albicans* to AMB, since AMB binds to ergosterol in the cell membrane^35^. Genes induced by both drugs in B11210 are enriched in amide biosynthetic process and translation, and genes only induced by VCZ are enriched in RNA processing and transcription (enriched GO terms corrected-*P <* 0.05, hypergeometric distribution with Bonferroni correction; **Table S9**).

Since *C. auris* isolates B8441 and B11210 (clade I) had disparate resistance phenotypes predominantly in response to AMB, we further examined expression changes between these two strains. Despite the fact that B8441 and B11210 are from the same clade (clade I) only 8 genes were induced in both strains upon AMB treatment, including *ARG1*, *CSA1* and *MET15*, and one *OPT1*-like transporter (B9J08_001998) (**Table S10**). In addition, comparison of both isolates before treatment revealed that the resistant strain B11210 has higher expression of genes previously noted to be involved in the *C. albicans* transcriptional response to drug exposure relative to B8441 even in the absence of drug (**Table S11**). This set of genes encompasses the D-xylulose reductase (*XYL2*)^35^, Phosphoenolpyruvate carboxykinase (*PCK1*)^26,35^, and a large set of transporters (*FRP1*, *FTH1*, *HGT13*, *HGT7*, *HSP70*, *NGT1*, *OPT1*, *PTR22*, *SEC26*)^35^, which suggests intrinsic expression of genes associated with polyene resistance in B11210 (**Table S10**). In addition, we observed that the expression levels of homologs of multidrug transporters (*CDR1*, *CDR4a*, *CDR4b*, *MDR1*, *SNQ2*) did not significantly vary across exposure to either drug or between isolates. Of these, *CDR4a* appears more highly transcribed relative to other multidrug transporters (*CDR1*, *MDR1* or *SNQ2*). This supports previous findings of intrinsic higher expression of transporters as a possible mechanism of drug resistance in *C. auris*^12^.

We compared these results to SNP variants identified between B8441 and B11210 to evaluate if any may explain the phenotypic difference in drug resistance in these strains. We found a total of 1,148 SNPs and 430 insertion or deletion events (**Table S11**). A total of 15 genes display a loss of function mutation (frame-shift, stop gained or stop lost), including the stationary phase protein (SNZ1) and the putative GPI-anchored adhesin-like protein (*HYR3*). We did not find loss of function mutations in genes previously associated with amphotericin B resistance in *C. albicans*, however, we did identify non-synonymous mutations in genes associated with drug resistance and response to amphotericin B, including *ERG11*, *ERG4*, *CDR1, C5_05230C_A, NHP6A* and other transporters such as *HGT7* and *YCF1,* (**Table S11**). These variants suggest candidate mutations that could explain phenotypic variation in drug resistance, however a wider association study of mutations and drug resistance levels would help identify the major genes involved across the different clades.

While many genes associated with antifungal resistance are conserved across *C. auris* and CTG species, unique genes and gene duplication in *C. auris* may also contribute to the underlying emergence of multidrug resistant phenotypes. Some genes induced by AMB or VCZ in *C. auris* B8441 and B11210 (**Table S10**) were specific to *C. auris* or to the emerging clade of *C. auris*, *C. haemulonii*, *C. duobushaemulonii,* and *C. pseudohaemulonii*. This includes five ortholog families unique to *C. auris*, comprising three putative GPI-anchored cell wall proteins, two homologs of *IFF6* (a putative GPI-anchored adhesin-like protein), one homolog of *PGA54*, and an aspartyl protease similar to *SAP8* (**Table S9**). In addition, this set included gene families that are expanded in *C. auris* or in closely related species (**Figures 3b** and **4**), including transporters (four *SIT*-like, one *FTR*-like, and two *OPT*-like class), cell wall adhesins, and several predicted secreted proteins (**Table S9**). Notably, while other Saccharomycetales species including *C. albicans* only have one copy of *SIT1* that is induced during AMB treatment, *C. auris* and closely related species had up to 11 *SIT1*-like genes, of which 4 are induced during AMB treatment (**Figures 3b**, **4**, **S6)**. Together, these transcriptional changes highlighted shared and *C. auris*-specific genes that might contribute to the MDR phenotype observed in *C. auris* isolates and provided candidate genes to further investigate *C. auris* multidrug-resistance.

## Discussion

As an emerging pathogen, *Candida auris* has not been well studied to date, highlighting the need for rapidly closing this knowledge gap to respond to the increasing number of fatal infections. There is also a limitation on how much of the biology of *C. auris* we can infer from related *Candida* species; *C. auris* is distantly related to the two most commonly observed pathogenic *Candida* species, *C. albicans* and *C. glabrata*, as well as other sequenced species. Our comparative genomic analyses, incorporating new genomic data for more closely related, multidrug-resistant species, revealed the recent evolution of this group of emerging pathogens including shared properties that underlie antifungal resistance and virulence. In addition to *C. auris* clades, we generated annotated genomes for three other closely related species rarely observed as infecting humans. Building on prior studies of individual loci^4,5,12,13^, the phylogenetic relationship of these species was more clearly resolved by whole genome comparisons; *C. haemulonii*, *C*. *duobushaemulonii*, and *C. pseudohaemulonii* are more closely related to each other than to *C. auris*, with the closest relationship being between *C*. *duobushaemulonii*, and *C. pseudohaemulonii*. Our phylogenetic analysis integrating the genomes of these species with other *Candida* highlight the distant relationship of this group to other pathogenic *Candida* species and the placement of these species within the CTG clade.

To characterize mechanisms that may contribute to virulence and drug resistance, we compared the gene content between the emerging multidrug resistant species and other related *Candida*. Recent work found that virulence in *C. auris* appears similar to *C*. *albicans* and *C*. *glabrata*^36^, suggesting that shared gene content could play a role. *C. auris* shares some notable gene family expansions described in *C. albicans* and related pathogens^23^, including of transporters and secreted lipases. While an expansion of transporters is shared, species-specific expansions have contributed to the diversification of transporters in *C. auris* and closely related species. Similarly, the expansion of lipases suggests this could be part of a shared mechanism of virulence; however, the roles of specific genes will need to be investigated. In contrast, expansions of cell wall families detected in *C. albicans* and related pathogens are not found in *C. auris* and related species. For example, the *ALS* family is represented by two to four copies in *C. auris* and related MDR species, compared to the eight copies present in *C. albicans*. Examining the predicted cell wall proteins in *C. auris* did not reveal any highly expanded families; perhaps the longer history of association with humans has impacted diversification of such cell wall protein families in the more commonly observed pathogenic *Candida* species.

Drug resistance in *C. auris* likely involves mechanisms previously described in *C. albicans*, however the specific transporters involved in drug response are less well conserved. All four species are resistant to antifungal drugs, and the Y132F mutation in the azole drug target *ERG11*, previously described in *C. auris*, and was also detected in *C. pseudohaemulonii*. The direct link between these mutations and azole resistance in *C. auris* is supported by recent work that found that expression of either the Y132F and K143R *C. auris ERG11* allele led to an increase in azole resistance in a heterologous system^17^. In addition, we find evidence of increased copy number of *ERG11* in two *C. auris* isolates, but little evidence of multiple isolates with regions of copy number variation, and no evidence of aneuploidy. This suggests that increased copy number of *ERG11* may be a mechanism of drug resistance in *C. auris*, as has been described in *C. albicans*^37^. However other mechanisms of drug resistance may vary between species and strains. For example, we found that efflux proteins such as the *CDR* family were not induced during voriconazole treatment, instead other transporters were up-regulated, including those that are either expanded or unique in *C. auris* and closely multidrug emergent species, highlighting that different molecular mechanisms are likely involved in the drug response.

Our analysis identified representative *C. auris* isolates for each of the two mating types, suggesting that this species has the potential to undergo mating and meiosis. Each of the four described *C. auris* clades consists of isolates that are all *MTL**a*** or *MTL*α; as the clades are geographically restricted, this suggests that there is a geographic barrier to opposite sex mating. This expectation would change if wider sampling demonstrated the establishment of both mating types within a geographic area. The potential for mating within this species is supported by the conservation of genes involved in mating and meiosis; these patterns are similar to that of the related species *C. lusitaniae*, for which mating and recombination have been demonstrated^24^. Isolates from clades of opposite mating types could be directly tested for mating and production of progeny. One potential barrier to mating between clades is the presence of chromosomal rearrangements, as some recombination events may result in inviable progeny.

Analyses of these genomes have revealed fundamental aspects of these emerging multidrug resistant fungi. The role of specific genes in mating, drug resistance or pathogenesis needs to be directly tested, utilizing gene deletion technologies recently adapted for *C. auris* (e.g. ^38^). Further analysis of this data will not only advance our understanding of the basis of drug resistance and virulence of this pathogen but can also inform development of fungal diagnostics for accurate tracking of these emerging pathogens.

## Methods

### Selected isolates and genome sequencing

Strains used in this study were described previously^2,18,19^; brief details are presented in **Table 1**. Isolates were grown on Sabouard Dextrose media supplemented with chloramphenicol and gentamycin and incubated for 24-48 hours at 37°C. For Illumina sequencing, genomic DNA was extracted using the *Quick*-DNA™ (ZR) Fungal/Bacterial Miniprep Kit (Zymo Research, Irvine, CA, USA). Genomic libraries were constructed and barcoded using the NEBNext Ultra DNA Library Prep kit (New England Biolabs, Ipswich, MA, USA) by following manufacturer’s instructions. Genomic libraries were sequenced using either Illumina HiSeq 2500 with HiSeq Rapid SBS Kit v2 or Illumina MiSeq platform using MiSeq Reagent Kit v2 (Illumina, San Diego, CA, USA). For PacBio sequencing, DNA was extracted using MasterPure™ Yeast DNA Purification Kit (Epicenter, Madison, WI, USA). Single-molecule real-time (SMRT) sequencing was done using the PacBio RS II SMRT DNA sequencing system (Pacific Biosciences, Menlo Park, CA, USA). Specifically, 20-kb libraries were generated with the SMRTbell Template Prep Kit 1.0 (Pacific Biosciences). Libraries were bound to polymerase using the DNA/Polymerase Binding Kit P6v2 (Pacific Biosciences), loaded on two SMRTcells (Pacific Biosciences), and sequenced with C4v2 chemistry (Pacific Biosciences) for 360 min movies.

### Genome assemblies

The genomes of B8441 and B11221 *C. auris* isolates were assembled as described^2^. *C*. *haemulonii* and *C*. *duobushaemulonii* genomes were assembled using Canu v1.6^39^. The resultant contigs were checked for further joins and circularity using Circlator v1.5^40^. The final contigs were polished using Quiver, part of SmrtAnalysis suite v2.3 (Pacific Biosciences)^41^. The sequence order for the chromosomes was verified using restriction enzyme AflII Whole Genome Mapping (OpGen, Gaithersburg, MA).

The sequenced Illumina reads of *C. auris* (B11220 and B11243), and *C. pseudohaemulonii* (strain B12108) were assembled using the SPAdes assembler v3.1.1^42^. Next, Pilon v1.16^43^ was used to polish the best assembly of each isolate, resolving single nucleotide errors (SNPs), artifactual indels and local mis-assemblies. All genome assemblies were evaluated using the GAEMR package (http://software.broadinstitute.org/software/gaemr/), which revealed no aberrant regions of coverage, GC content or contigs with sequence similarity suggestive of contamination. Scaffolds representing the mitochondrial genome were separated out from the nuclear assembly. All genome assemblies have been deposited at deposited at DDBJ/EMBL/GenBank (see ***Data availability statement***). To address if any strain representing each *C. auris* clade could be uniformly diploid, we examined candidate heterozygous positions predicted by Pilon v1.12^43^ using mapped Illumina data. The low frequency and absence of such positions supported that all sequenced genomes are homozygous haploids.

### Optical mapping

Two strains of *C. auris* (B11221 and B8441) were compared by the OpGen optical mapping platform (OpGen, Inc., Gaithersburg, Maryland). High molecular weight genomic DNA from overnight grown cells were purified with Argus HMW DNA Isolation Kit (OpGen, Inc.) and examined for quality and concentration using the ARGUS QCards(OpGen, Inc.). The software program Enzyme Chooser (OpGen, Inc.) identified *BamHI* restriction endonuclease to be optimal for optical map production, because its cleavage of reference genomes would result in fragments that average 6–12 kbp in size, with no fragments larger than 80 kbp. Single genomic DNA fragments were loaded onto a glass surface of a MapCard (OpGen, Inc.) using the microfluidic device, washed and then digested with *BamHI* restriction enzyme, and stained with JOJO-1 dye through the ARGUS MapCard Processor (OpGen, Inc.). Map cards were scanned and analyzed by automated fluorescent microscopy using the ARGUS Whole Genome Mapper (OpGen, Inc.). The single molecule restriction map collections were then tiled according to overlapping fragment patterns to produce a consensus whole genome map. This map was imported into MapSolver (OpGen, Inc.) along with predicted *in silico* maps of contigs derived from WGS, using the same restriction enzyme for ordering and orientation of contigs during genome circularization. *In-silico* predicted optical maps of complete genomes were scaled according to the size of sequenced genomes to show identity with Optical maps (**Figures S1**).

### RNA-Seq of B8441 and B11210 during drug treatment

For RNA extraction *C. auris* cells were grown in YPD broth medium (Difco Laboratories, Sparks, MD) at 30°C in a shaking incubator at 300 rpm. After 18 hours, the stationary phase cells were diluted with the equal volume of fresh YPD broth and incubated for two hours at 37° to induce growth. After that, the cells were treated with amphotericin B at the final concentration of 0.25 µg/mL or voriconazole at the final concentration of 1 µg/mL (Sigma-Aldrich, St. Lois, MO) and incubated at 30°C for additional 2 and 4 hours in the presence of drug. Cells were centrifuged for 2 minutes at 12000xg, pellets were flash frozen in dry ice/ethanol bath and stored at -80°C. RNA was isolated using RiboPure™-Yeast rapid RNA isolation kit (Life technologies, Carlsbad, CA) using the manufacturer’s protocol. RNA was adapted for sequencing using the RNAtag-Seq approach^44^, with the modification that the yeast RiboZero reagent was used for rRNA depletion. For each condition, two biological replicates were performed, and the read counts per transcript were highly correlated between replicates (R> 0.90).

### Gene annotation

Gene annotation in *C. auris* was performed using RNA-Seq paired-end reads to improve gene calling and structure predictions. Briefly, we mapped RNA-Seq reads to the genome assembly using Tophat2, and use the alignments to predict genes using BRAKER1^45^, that combines GeneMark-ET^46^ and AUGUSTUS^47^, incorporating RNA-Seq data into unsupervised training and subsequently generates *ab initio* gene predictions. Additionally, we re-annotated the genome of the 6684 strain^13^ improving its gene set and predicted gene structures. tRNAs were predicted using tRNAscan^48^ and rRNAs predicted using RNAmmer^49^. Genes containing PFAM domains found in repetitive elements or overlapping tRNA/rRNA features were removed. Genes were named and numbered sequentially. For the protein coding-gene name assignment we combined HMMER PFAM/TIGRFAM, Swissprot and Kegg products. For comparative analysis genes were functionally annotated by assigning PFAM domains, GO terms, and KEGG classification. HMMER3^50^ was used to identify PFAM domains using release 27. GO terms were assigned using Blast2GO^51^, with a minimum e-value of 1×10^-10^. Protein kinases were identified using Kinannote^52^ and transporter families using TCDB version 01-05-2017^53^. To evaluate the completeness of predicted gene sets, the representation of core eukaryotic genes was analyzed using CEGMA genes^54^ and BUSCO^55^.

### Copy number variation and nucleotide diversity

To identify regions in *C. auris* that exhibit copy number variation (CNV) we analyzed Illumina read depth of 47 previously sequenced isolates. We identified genomic windows (1 kb) showing significant variation (*p*-value < 0.01) in normalized read depth using CNVnator v0.3^56^. To measure nucleotide diversity we used SNPs identified from the isolate sequences of Lockhart *et al*^2^. We computed genome-wide π using VCFtools v0.1.12^57^ for non-overlapping sliding windows of 5 kb for *C. auris* clade I, II, II and all isolates.

### Comparative genomics and phylogenomic analysis

To examine the phylogenetic relationship of the emerging multidrug-resistance clade, including *C. auris*, *C. haemulonii*, *C. duobushaemulonii,* and *C. pseudohaemulonii* we identified single copy orthologs in these sequenced genomes and twelve related species using OrthoMCL v1.4^58^ (Markov index 1.5; maximum e-value 1e-5). A total of 1,570 protein-coding genes that have one single copy and are conserved in all 20 genomes were aligned using MUSCLE, and a phylogeny was estimated from the concatenated alignments using RAxML v7.7.8^59^ with model PROTCATWAG with a total of 1,000 bootstrap replicates. To compare gene family expansion and contractions, we used orthologous gene clusters we classified as core, auxiliary and unique. We then searched for expansions or contractions in functionally classified genes by assigning PFAM domains, GO terms, and KEGG classification. Using a matrix of gene class counts for each classification type, we identified enrichment comparing the emerging multidrug-resistance clade with all the other related species using Fisher’s exact test. Fisher’s exact test was used to detect enrichment of PFAM, KEGG, GO terms, and transporter families between groups of interest, and p-values were corrected for multiple comparisons^60^. Significant (corrected *P*-value < 0.05) gene class expansions or depletions were examined for different comparisons.

### Transcriptional analysis of *C. auris* RNA-Seq

RNA-Seq reads were aligned to the transcript sequences of *C. auris* B8441 or B11221 using Bowtie2^61^. Transcript abundance was estimated using RSEM (RNA-Seq by expectation maximization; v.1.2.21) as transcripts per million (TPM). TPM-normalized ‘transcripts per million transcripts’ (TPM) for each transcript were calculated, and differentially expressed transcripts were identified using edgeR^62^, all as implemented in the Trinity package version 2.1.1^63^. Genes were considered differentially expressed only if they had a 2-fold change difference (> 2 FC) in TPM values and a false discovery rate below or equal to 0.05 (FDR < 0.05), unless specified otherwise. To determine major patterns of antifungal-response specific we clustered gene expression patterns by *k-means*. To identify functional enrichment of differentially expressed genes, we used functional gene assignations from PFAM, GO terms and KEGG (see ***Gene annotation***), and then performed comparisons with Fisher’s exact test. To identify possible functions of the gene products of significantly differentially expressed drug resistance genes, protein homologs were assigned based on orthology, functional assignments (GO, PFAM, TIGRFAM) and experimental evidence from *Candida* genome database (http://www.candidagenome.org).

### Antifungal susceptibility testing

Antifungal susceptibility testing was performed on isolates according to Clinical and Laboratory Standards Institute (CLSI) guidelines^64^. Custom prepared microdilution plates (Trek Diagnostics, Oakwood Village, OH, USA) were used for generating both azole and echinocandin susceptibilities while Etest® strips (BioMerieux, Marcy l’Etoile, France) were used for amphotericin B susceptibilities. Interpretive breakpoints for *C. auris* and related species do not exist but have been conservatively based on those breakpoints established for other *Candida* species.

### Data availability statement

All genome assemblies and gene annotations have been deposited at DDBJ/EMBL/GenBank under the following accession numbers: *C. auris* B8441 PEKT00000000; *C. auris* B11221 PGLS00000000, *C. auris* B11220 PYFR00000000, *C. auris* B11243 PYGM00000000, *C. haemulonii* B11899 PKFO00000000, *C. duobushaemulonii* B09383 PKFP00000000, *C. pseudohaemulonii* B12108 PYFQ00000000. All whole genome sequence data have been deposited in NCBI under the following BioProjects: *C. auris* PRJNA328792, *C. haemulonii* PRJNA421961, *C. duobushaemulonii* PRJNA421966, *C. pseudohaemulonii* PRJNA438484.The RNA-Seq data from *C. auris* has been deposited at NCBI under BioProject PRJNA445471.

## Acknowledgements

We thank Sarah Young and Paul Cao for their assistance with genome annotation, the Broad Technology labs for RNAtag-Seq library construction, and the Broad Genomics Platform for Illumina sequencing. This project has been funded in whole or in part with Federal funds from the National Institute of Allergy and Infectious Diseases, National Institutes of Health, Department of Health and Human Services, under award N°: U19AI110818. The use of product names in this manuscript does not imply their endorsement by the US Department of Health and Human Services.

## Disclaimer

The finding and conclusions in this article are those of the authors and do not necessarily represent the views of the CDC.

## Author contributions

Conceived and designed the study: APL CAC. Performed experiments: LG NAC VNL ELB PJ. Performed the assembly and annotation: JFM. Analyzed the data: JFM VNL PJ RAF CAC. Wrote the paper: JFM CAC.

## Additional information

Competing interests: The authors declare no competing interests.

